# The architecture and design of ecological null models

**DOI:** 10.1101/195131

**Authors:** Joshua Ladau

## Abstract

Many questions in ecology are best addressed using observational data because they concern spatial or temporal scales where experimentation is impractical. Null models, which make predictions in the absence of a particular ecological mechanism, are instrumental for making inferences in these situations, but which null models to use or how to best test them is often unclear; this ambiguity is problematic because different null models and tests can yield different results, suggesting contradictory ecological mechanisms. To address these challenges, this paper presents an overarching framework for the development and testing of null models, in which desirable models and tests are obtained as solutions to mathematical optimization problems. As an example of how the framework can be applied, this paper shows how it can be used to design null model tests to check for effects of interspecific interactions on species co-occurrence patterns. A minimal sufficient statistic (metric) for effects of interspecific interactions is derived, which achieves the maximal level of data compression without losing information present in the data about interspecific interactions. Existing, intuitive statistics are shown to lack this property. The paper then derives a statistical hypothesis test that has the greatest possible power (sensitivity) for detecting effects of competition and facilitation given a controlled false positive rate. This test is shown numerically to improve greatly over existing tests. The optimization paradigm allows the most accurate inferences possible, and should be applicable throughout ecology where null models are used to make inferences.

## 1 Introduction

Because of the large spatial and temporal scales at which ecological phenomena can occur, many key ecological questions must be addressed using non experimental, observational data (e.g., Connor and Simberloff, 1978, McNab, 1971). For instance, experimentally manipulating distributions of threatened species may be unethical, and it may be impossible to experimentally evaluate the effects of interspecific interactions on community structure at regional scales over hundreds of years (Connor and Simberloff, 1986). Financial limitations may also make experimentation possible only on a small number of ecological systems (e.g., Mitchell-Olds, 2001), while observational data can often be collected more readily, making them important for extrapolating from experimental results.

Statistical methods constitute a central means for making inferences about ecological processes from non-experimental data: they allow the consistency between patterns predicted from processes and patterns observed in data to be assessed. The statistical methods that are used in ecology fall into two categories: methods that have been developed in a general, non-ecological context (e.g., ANOVAs, Bayesian Information Criteria)(Sokal and Rohlf, 1995) and methods that have been developed specifically in the context of ecological questions (Gotelli and Graves, 1996). Methods of the latter kind have been created for situations not addressed by standard statistical methods, for instance, inference in models specific to ecology. They comprise a substantial proportion of the methods that are used in ecology (Gotelli, 2001, Dufrene and Legendre, 1997, Graves and Gotelli, 1993, Schluter, 1990, Gotelli and Abele, 1982).

This paper focuses on a class of methods developed specifically in ecological contexts: null model tests, wherein the consistency of data to a null model which posits the absence of a particular ecological process, is checked. Finding data inconsistent with the null model can suggest effects of that process. This paper will focus particularly on null model tests of presence-absence data (“NMTPAs”) (Connor and Simberloff, 1979), but the issues and approach developed here pertain directly to null model tests throughout ecology. Hence, in addition to being directly useful for the analysis of presence-absence data, the results presented here demonstrate a general approach with potentially broad applications.

NMTPAs are used to assess the effects of habitat filtering, competition, facilitation,and other interspecific interactions on large-scale distributions of species (Gotelli and Ellison, 2002, Dayton and Fitzgerald, 2001, Gotelli et al, 1997). In the last 25 years, they have been the used in over 250 published studies, including studies developing strategies for conservation (Feeley, 2003, Atmar and Patterson, 1993) and infectious disease control (Stephens et al, 2008). NMTPAs are currently in widespread use (Heino and Soininen, 2005, Mouillot et al, 2005, Gotelli and McCabe, 2002, Gotelli and Rohde, 2002), and their importance in community ecology is high (Sanders et al, 2003, Gotelli, 2001).

NMTPAs are statistical hypothesis tests. As inputs they use lists of species observed at sites or communities. These data are usually summarized in a presence absence ma trix, in which rows represent species and columns represent sites. If species *i* was observed at site *j*, then entry *i*,*j* of the presence absence matrix is 1; otherwise it is 0. As with all statistical tests, NMTPAs are rules that prescribe for every possible presence absence matrix one of two actions: *reject the null hypothesis* or *fail to reject the null hypothesis*. Equivalently, they can be regarded as methods for estimating confidence limits forpara meters of interest. The null hypothesis for NMTPAs has been defined in several ways, including that species colonize independently, they colonize “randomly,” they occur with equal probability at all sites, and they are equally likely to occur at different sites (Simberloff and Connor, 1981). These null hypotheses can be formalized mathematically, and inferences about them allow conclusions about ecological phenomena; for instance, if species colonize non independently (at Site I, species *A* is more likely to occur when species *B* is absent than when it is present), then effects of interspecific interactions might be suggested.

NMTPAs differ from standard frequentist statistical tests in one key aspect. Most standard statistical tests are solutions to optimization problems. For example, under certain assumptions, many standard tests (such as *t*-tests), give true positive results at the highest rate possible (i.e., have the greatest possible *power*), subject to giving false negative results at or below a specified rate (e.g., 0.05). Other tests may optimize other properties, optimize power subject to other constraints, or provide approximately optimal solutions. In contrast, NMTPAs have been developed largely on intuitive grounds, and optimality has not been explicitly used as a guide in their construction (e.g., Gotelli and Graves, 1996, Roberts and Stone, 1990, Wilson, 1989, Gilpin and Diamond, 1982, Connor and Simberloff, 1979).

There are at least two strong motivations for using optimality as a framework for constructing NMTPAs. First, an optimal test may be much more powerful (sensitive) than an intuitively based test. For instance, optimal tests may detect effects of interspecific interactions at over twice the rate of intuitive tests (see below). Thus, using optimal tests can lead to more accurate conclusions about the effects of ecological processes. The second motivation for optimal tests is the common situation that numerous NMTPAs can all seem intuitively reasonable for addressing a question of interest, but yield strikingly different conclusions when applied to the same data. Such problems have led to protracted controversy over which NMTPAs to use when (e.g, Brown et al, 2002, Stone et al, 2000, 1996, Fox and Brown, 1995, 1993), and indeed over the utility of null models tests in general (Gotelli and Graves, 1996, Alatalo, 1982). Optimality considerations can resolve such controversies.

This paper addresses two aspects of the optimality problem. The first regards the statistic (metric) that is used to measure effects of interspecific interactions. Statistics calculated from the data are used for summarizing co-occurrence patterns and constructing NMTPAs. However, some statistics capture more information about a process than others. This notion is formalized by the *sufficiency principle*: heuristically, if a statistic is sufficient, then it captures all of the available information. This paper derives a sufficient statistic for effects of interspecific interactions in the context of null models of presence absence data.

The second aspect of optimality that this paper addresses regards hypothesis testing. Using the sufficient statistic, the paper derives two *uniformly most powerful* (“UMP”) tests. Under certain conditions, these tests have the greatest power possible for any NMTPA that can be created (Lehmann and Romano, 2005, Casella and Berger, 2002). The properties of these NMTPAs are confirmed numerically, and the tests are applied to two ecological data sets. The use of these NMTPAs and the further development of optimal methods offers a promising new approach for addressing important unsolved inferential problems in ecology.

## 2 Analytic Results

This section illustrates how null model tests can be developed in a rigorous framework of optimization.

### 2.1 Model

The optimality of any statistical method (e.g., sufficiency of a statistic, UMP properties) depends greatly on what can be assumed about the process generating the data. For instance, if the data are known to be an independent sample from a normal distribution with known variance, then a *z*-test is UMP for testing whether the mean of the distribution is greater than a particular value. In contrast, if the sample is from a normal distribution with unknown variance, then a Student's *t*-test is UMP (Casella and Berger, 2002). This section describes the model of the generating process under which optimality results are derived here. We suggest that this model is reasonable for many circumstances. In other circumstances, other mossdels may be more appropriate, and the resultant optimal NMTPAs may be different.

The model, denoted ℳ, is characterized by four assumptions that are defined precisely in section 6.1. Heuristically, the first assumption is that communities assemble independently of each other; e.g., the species occurring at Site I do not affect which species occur at Site II. This assumption is common in NMTPAs and is likely to be largely justified for many data sets where dispersal is limited. The second assumption regards the equivalence of species: unconditional on the presence of other species, all communities with the same numbers of species are equally likely to occur. For data sets concerning ecologically similar species, this assumption appears reasonable. Moreover, it is consistent with neutral theory (Hubbell, 2001), which posits extensive equivalence between species.

The last two assumptions regard species equivalence and the time scale of community assembly. In the presence of antagonistic interspecific interactions, the assumptions posit that unconditional on the presence of other species, each species occurs with probability *p*. However, conditional on the presence of one or more species, the probability drops to *pξ*, where *ξ <* 1. Hence, antagonistic interactions have the same effect regardless of whether one or many species are present in a community. Facilitation has corresponding effects, with *ξ >* 1. This scenario is consistent with the following ecological process: Suppose that species are competitively equivalent, and that new species arrive at a community at a low rate. Thus, when the first species arrives, its population will reach the carrying capacity of the community before the second species arrives. When that species arrives, it will encounter a “full” community, and hence have a depressed probability of successfully colonizing with antagonistic interactions, or an increased probability with facilitation. If it does colonize, then the community will again have time reach equilibrium, and when the third species arrives, it will encounter a “full” community. By the equivalence of species,the third species will have the same reduced or increased probability of occurring as the second species. The process iterates. Such a scenario appears particularly reasonable if communities are spatially isolated, and it accords with the equivalence postulated by neutral theory (Hubbell, 2001).

### 2.2 Sufficient Statistics

In the context of making a statistical inference, a statistic is means for compressing data. However, certain statistics compress data better than others. For instance, suppose that we wish to determine the bias of a coin, and to do so we flip it 5 times, obtaining the sequence of outcomes *HTTHH*. Intuitively, this sequence contains information that is both relevant and irrelevant to assessing the bias. The total number of heads is clearly relevant, but the exact order in which they occurred is irrelevant no additional informa tion about the bias is gained by knowing that *HTTHH* rather than *THTHH* occurred. Hence, a statistic that depended only on the ordering would be undesirable, one that depended only on the number of heads would be desirable, and one that depended on both would be less desirable, having failed to compress the data completely. Heuristically, a *sufficient statistic* for a model does not lose information about that model, and *minimal sufficient statistic* achieves the greatest degree of data compression possible without losing information (Casella and Berger, 2002, Lehmann and Casella, 1998).

Let the statistic *U*_0_ give the number of sites in a presence-absence matrix at which no species were observed, *S* give the total number of species-occurrences in the matrix, and **T**_*U*_ be the ordered pair (*U*_0_, *S*). Theorem 1 follows:

**Theorem 1.** *The statistic* **T**_*U*_ *is minimal sufficient for ℳ.*

As noted above, numerous statistics have been proposed for summarizing co-occurrence patterns. It is reasonable to ask whether these statistics are also sufficient for *ℳ*. Four widely considered statistics are the *Number of Checkerboards*, *C-Score*, *Number of Species Combinations*, and *V-Ratio* (Gotelli, 2000). The Number of Checkerboards gives the number of species having non overlapping distributions in a presence absence matrix (Diamond, 1975). The *C* Score is the number of “checkerboard units” (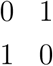 and 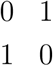 submatrices) in a presence absence matrix divided by the number of pairs of species (Stone and Roberts, 1990). The Number of Species Combinations is the number of unique assemblages of species, where assemblages are considered on a per site basis (Pielou and Pielou,1968). The *V-*ratio is a ratio of variances, calculated using the marginal totals of the presence absence matrix (Schluter, 1984). Let *T*_*Ch*_,*T*_*C*_,*T*_*Co*_, and *T*_*V*_ denote these statis tics, respectively. Although all of the statistics can be justified intuitively, the following theorem holds:

**Theorem 2.** *The statistic T*_*C*_, *T*_*Ch*_, *T*_*Co*_, *and T*_*V*_ *are not sufficient for ℳ.*

### 2.3 Null Hypothesis

The optimality of any hypothesis test depends on the conclusions that are drawn from it; the null hypothesis that it is used to check. A test that yields highly reliable conclusions about one null hypothesis may yield unreliable conclusions about another null hypothesis. In this paper, two tests are developed, a “Competition Test” and a “Facilitation Test.” These tests are optimal for different null hypotheses. It follows from the definition of *ξ* that the null and alternative hypotheses for testing for antagonistic interspecific interactions (e.g., competition) are

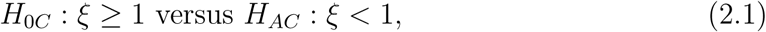

and the hypotheses for testing for facilitation are

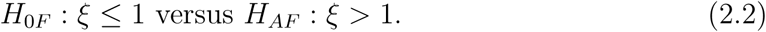

Here, “competition” and “facilitation” are taken mean negative and positive non independence in community assembly, respectively. They can suggest effects of negative and positive interspecific interactions, but additional lines of evidence would of course be necessary to definitively conclude effects of such interactions from observational data.

### 2.4 Uniformly Most Powerful Tests

This section presents UMP tests of *H*_0*C*_ versus *H*_*AC*_, and *H*_0*F*_ versus *H*_*AF*_. The UMP tests make use of the sufficient statistic **T**_*U*_, derived above. In general, sufficient statistics are crucial for creating optimal hypothesis tests because heuristically, inferences made using statistics that discard pertinent information are less reliable than those made using all of the available information. The UMP tests have two additional useful properties.First, they can be of any specified size *α* ∊ [0, 1]; i.e., they give false positives at a rate not exceeding *α*. Second, the UMP tests are unbiased, always rejecting the null hypothesis more often when it is true than when it is false. This makes them UMPU tests (Casella and Berger, 2002).

The UMPU tests are randomized tests; i.e., for any presence absence matrix **b**, they reject *H*_0_ with probability *ϕ*(**b**). The motivation for randomization is to allow tests with an exact size (e.g., 0.05) to be created, which is otherwise impossible because of the discrete distributions involved. The function *ϕ* is called the critical function, and it completely characterizes a randomized test. Here, *ϕ* will also be used to refer to the test itself. Let *m* and *n* be the number of rows and columns in the observed presence absence matrix, respectively, and let Ω be the set of all possible *m × n* presence absence matrices. Define *P*_0_ as the distribution of presence absence matrices wherein every matrix in Ω is equally likely. The following general definition will be useful:

**Definition 1.** *For any* **b** ∊ Ω, *statistics T*_1_ *and T*_2_ *on* Ω, *and k ∊ {−*1, 1*}, let*

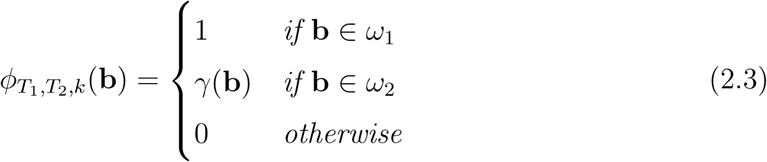

*where ω*_1_, *ω*_2_, *and γ are defined as follows:*

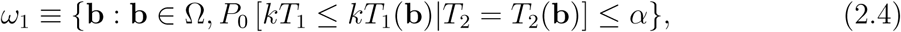

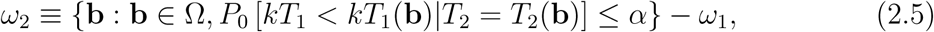

*and γ* : Ω *→* [0, 1] *such that*

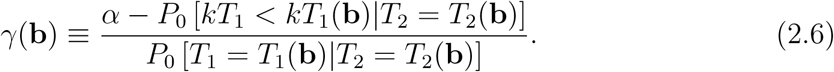

Theorems 3 and 4 present the UMPU tests of *H*_0*C*_ against *H*_*AC*_ and *H*_0*F*_ against *H*_*AF*_, respectively:

**Theorem 3.** *ϕ*_*U*0, *S,*1_ *is a UMPU size-α test of H*_0*C*_ *versus H*_*AC*_.

**Theorem 4.** *ϕ*_*U*0, *S,−*1_ *is a UMPU size-α test of H*_0*C*_ *versus H*_*AC*_.

For convenience, these tests will be referred to as *ϕ*_*UC*_ (“UMPU Competition”) and *ϕ*_*UF*_ (“UMPU Facilitation”), respectively. Both of these tests may be difficult to implement because of difficulties in calculating *P*_0_ exactly. However, good numerical approximations are available, which are described in the next section.

## 3 Numerical Methods

This section shows how the optimal tests derived above can be implemented, and characterizes their behavior numerically.

### 3.1 Numerical Approximations to *ϕ*_*UC*_ and *ϕ*_*UF*_

The following is a procedure to approximate *ϕ*_*UC*_ numerically for an an observed presence-absence matrix **b**^*\**^:

1. Simulate *P*_0_: Sample equiprobably from the set of presence absence matrices with *m* rows, *n* columns, and *S*(**b**^*\**^) species occurrences.
2. Find the corresponding simulated distribution of *U*_0_: For each simulated matrix, the value of *U*_0_ is the number of columns containing no species.
3. If it exists in the simulated distribution, find the maximum value *u*_0_^*\**^ such that 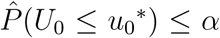. Otherwise, set *u*_0_^*\**^ equal to the minimum simulated value of *U*_0_ minus 1.
4. In the simulated distribution, find the next largest value to *u*_0_^*\**^, here denoted *u*_0_^*\*\**^.
5. if *U*_0_(**b**^*\**^) *≤ u*_0_^*\**^, reject *H*_0*C*_. If *U*_0_(**b**^*\**^) *> u*_0_^*\*\**^, then fail to reject *H*_0*C*_.
6. If *U*_0_(**b**^*\**^) = *u*_0_^*\*\**^, then compute

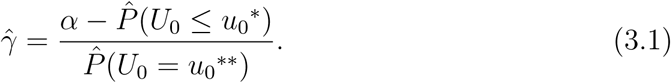

If a uniform [0, 1] random deviate is less than *γ*, reject *H*_0*C*_; otherwise fail to reject it.

To approximate *ϕ*_*UF*_, the same steps can be followed, except the inequality signs (excluding “*≤ α*”) are reversed and *u*_0_^*\**^ is set equal to the maximum simulated value of *U*_0_ plus 1 in the second part of step 3. To simulate *P*_0_ in step 1, a Markov Chain Monte Carlo (MCMC) algorithm (Ross, 2006) may be used. An implementation using the Metropolis Hastings algorithm is as follows:

1. Simulate an initial presence absence matrix with *S*(**b**^*\**^) species occurrences: Ran domly select *S*(**b**^*\**^) distinct numbers from { 1, *…, mn}*. Cell (*i, j*) contains a 1 in the presence absence matrix if (*i −* 1)*n* + *j* is among these numbers; it is 0 otherwise.
2. Randomly choose two cells in the matrix. If the values of the cells differ, then swap them; otherwise do nothing. Save the resulting matrix, even if no swap was performed.
3. Iterate step 2.

The list of matrices from step 2 will closely approximate *P*_0_ when a large number of iterations are performed. The speed of the algorithm makes it possible to perform *≈* 10^5^ iterations.

### 3.2 Power Functions

Section 6.2 proves that *ϕ*_*UC*_ and *ϕ*_*UF*_ are UMPU for the corresponding null and alternative hypotheses. To confirm these results and investigate how much *ϕ*_*UC*_ and *ϕ*_*UF*_ improve over existing NMTPAs, I numerically evaluated *power functions* for *ϕ*_*UC*_, *ϕ*_*UF*_, and 16 commonly used NMTPAs. A power function gives the probability of rejecting the null hypothesis over the possible values of *ξ*. Hence, for *ξ ∊ H*_0_, it gives the Type I error rate, and for *ξ ∊ H*_*A*_, it gives the power.

The eight NMTPAs that I used implemented the four statistics discussed in Section 2.2, and conditioned on the total number of species occurrences or the row/column totals. To define these tests, let the vector **M**(**b**) give the marginal totals of presence absence matrix **b**. I considered the following tests for effects of antagonistic interspecific interactions: 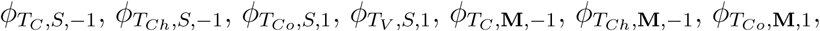 and 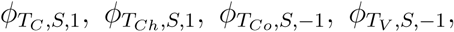. The corresponding tests for effects of facilitation were: 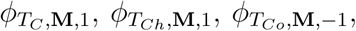 and 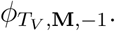. Thetests 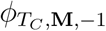 and 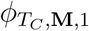 are currently in particularly widespread use; they implement the *C* score statistic and condition on the row and column totals (e.g., Gotelli and McCabe, 2002).

The power functions of these tests depend on three conditions. The first is the dimen sions of the presence absence matrix. The results reported here are for a 8 *×* 26 matrix of seabirds in the United Kingdom (Reed, 1980), which appears to yield generally representative results. The second condition is the value of the occurrence probability *p*, which is a nuisance parameter. Here, I set *p* = 0.25. The third condition is the generating process. As mentioned above, assumptions about the generating process can strongly affect the optimality of a statistical method. Hence, I considered three models:

1. *ℳ*: The model described above under which *ϕ*_*UC*_ and *ϕ*_*UF*_ were proven UMP unbiased.
2. *ℳ*_1_: A model equivalent to *ℳ*, except that the assumption of site equivalence is relaxed; species may have varying unconditional probabilities of occurring at different sites.
3. *ℳ*_2_: A model equivalent to *ℳ*, but with the assumption that effects of interspecific interactions occur after 1 species has colonized relaxed. The effects are assumed possible after an arbitrary number of colonizations.

To evaluate the power functions, I simulated 5000 presence absence matrices assuming the three models and different values of *ξ* ∊ [0.85, 1.6] (below *ξ ≈* 0.85 with *n* = 8, *ℳ* is self inconsistent). For *ℳ*_1_, I assumed that the probability of occurrence for each site was a sample from a uniform distribution on (0.15, 0.35). For *ℳ*_2_, I assumed that interspecific interactions could have effects after the fourth species had arrived. I checked whether each simulated matrix was in the critical region of each test, setting *α* = 0.05 and using randomized tests to allow fair comparisons. I implemented the tests using unbiased Metropolis Hastings algorithms with chain lengths of 10^5^ (Ross, 2006, Miklos and Podani, 2004). The overall rejection frequency per value of *ξ* provided an estimate of the power function at that value of *ξ*. Software for implementing these procedures was written in Visual Basic 6.0 and is available upon request.

### 3.3 Application

To illustrate the application of *ϕ*_*UC*_ and *ϕ*_*UF*_, and to compare the results of these tests to those of the NMTPAs listed in Section 3.2, I analyzed two published presence absence matrices. The first presence absence matrix was for United Kingdom seabirds (see section 3.2) (Reed, 1980). The second presence absence matrix was for ants in Virginia,with dimensions 11 *×* 25 (Gotelli, 2000). These presence absence matrices have been the subject of previous NMTPA analyses, and in addition, the authors were unbiased against reporting sites without species, a necessity for *ϕ*_*UC*_ and *ϕ*_*UF*_. Results are reported for both randomized and non randomized versions of each test.

## 4 Numerical Results

Figure 1 shows the power functions of *ϕ*_*UC*_, *ϕ*_*UF*_, and the other NMTPAs that condition on *S*. Under *ℳ*, the power of *ϕ*_*UC*_ and *ϕ*_*UF*_ greatly exceeded that of the other tests, being as high as 0.79 when the other tests were below 0.44.In addition, *ϕ*_*UC*_ and *ϕ*_*UF*_ had the lowest Type I error rates. For *ℳ*_1_, *ϕ*_*UC*_ and *ϕ*_*UF*_ still outperformed other tests, although the power of all of the tests was reduced and the size of several of the tests was close to 0.1. Under *ℳ*_2_, all of the tests lacked power, and some, including *ϕ*_*UC*_ and *ϕ*_*UF*_, were biased. The NMTPAs that conditioned on **M** almost universally lacked power, and under *ℳ* and *ℳ*_1_, *ϕ*_*UC*_ and *ϕ*_*UF*_ had much greater power and lower Type I error rates (Figure 2).

**Figure 1:**
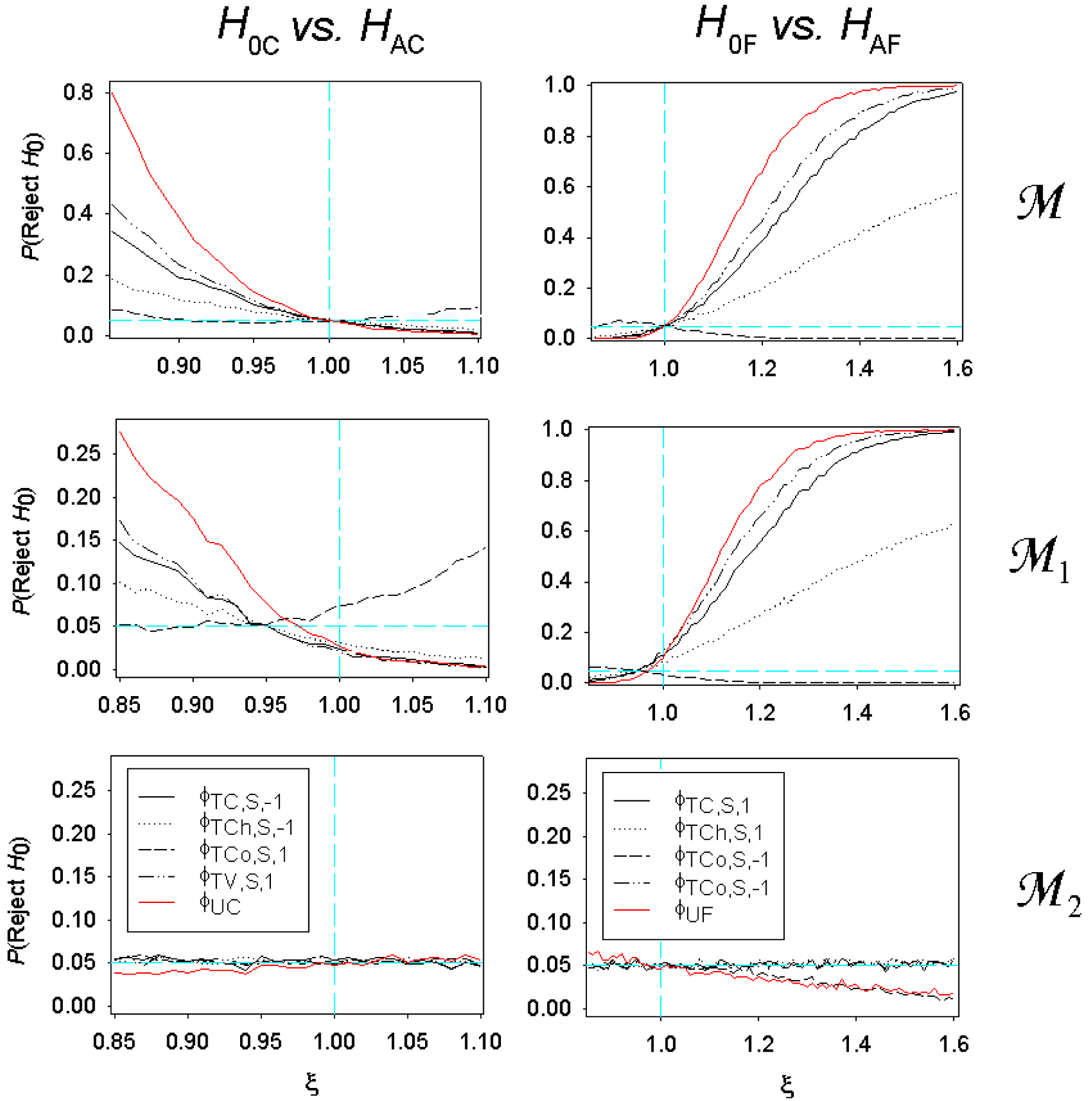
Power functions of *ϕ*_*UC*_, *ϕ*_*UF*_, and other NMTPAs conditioning on *S*. The first column shows power functions for *ϕ*_*UC*_ and other tests of *H*_0*C*_ (left legend). The second column shows power functions for *ϕ*_*UF*_ and tests of *H*_0*F*_ (right legend). The power functions in the first row are under model *ℳ*; here *ϕ*_*UC*_ and *ϕ*_*UF*_ can be proven UMP. The power functions in the second and third rows are under models *ℳ*_1_ and *ℳ*_2_.

**Figure 2:**
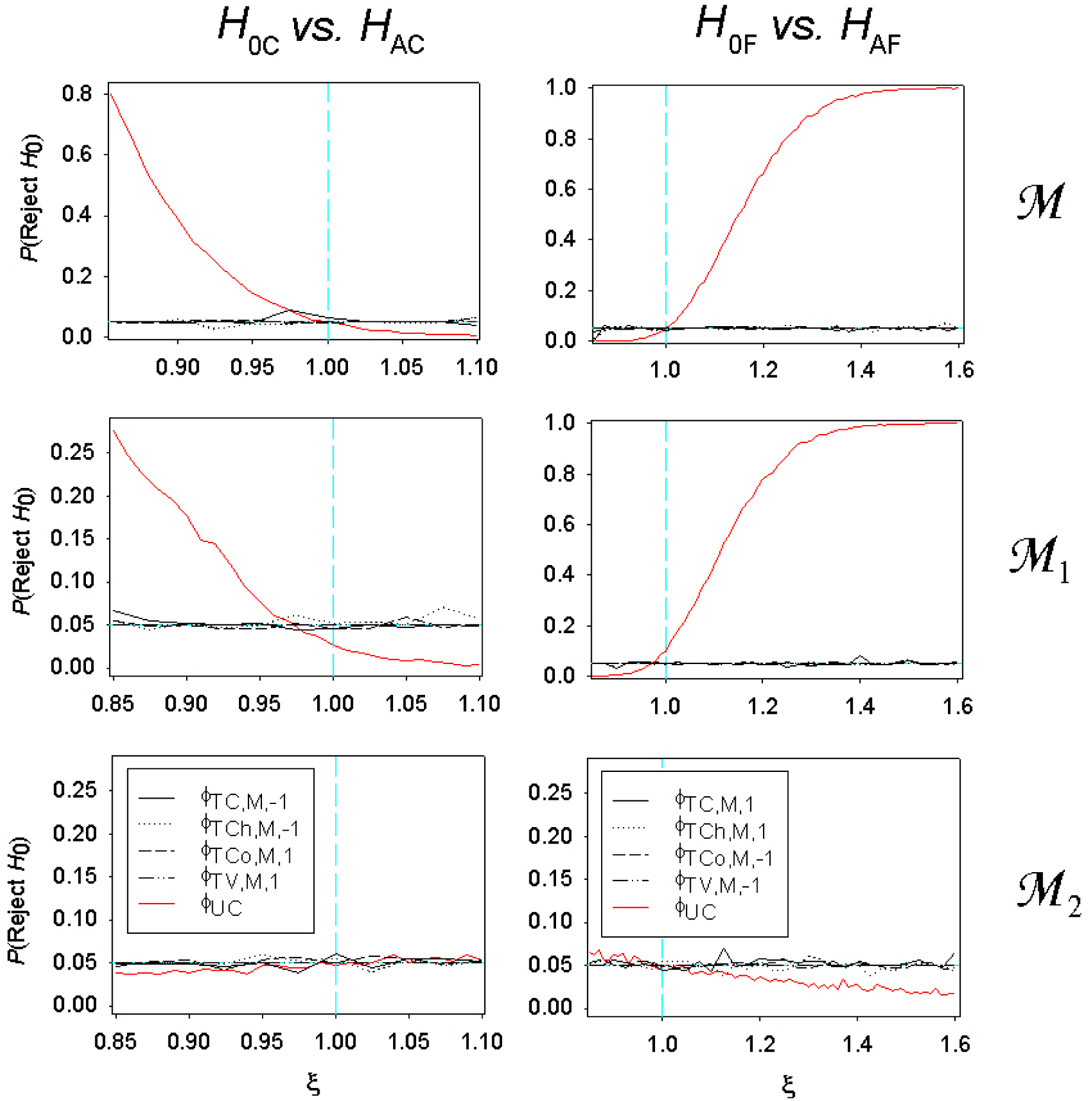
Power functions of *ϕ*_*UC*_, *ϕ*_*UF*_, and other NMTPAs conditioning on **M**. The plots are analogous to those in Figure 1.

The results of the analyses of the seabird and ant presence absence matrices are presented in Table 1. For the UK seabirds data, *ϕ*_*UC*_ and *ϕ*_*UF*_ never rejected the null hypothesis, although other tests did. This is likely due to model misspecification or the latter tests having higher Type I error rates. Similar results held for the Virginia ant data, with *ϕ*_*UC*_ and *ϕ*_*UF*_ never rejecting the null hypothesis. Overall, with one exception, the randomized and non-randomized versions of each test gave identical results.

**Table 1:** Results of applying the NMTPAs to the UK seabird and Virginia ant data sets. Tests rejecting the null hypothesis are listed, categorized by randomization and the null and alternative hypotheses that they tested. A list of all of the tests that were examined is given in section 3.2.

| Hypotheses | UK Seabirds |  | Virginia Ants |  |
| --- | --- | --- | --- | --- |
|  | Randomized | Nonrandomized | Randomized | Nonrandomized |
| $H_{0C}$ vs. $H_{AC}$ | $\phi_{T_C,S,-1}, \phi_{T_{Ch},S,-1}$ | $\phi_{T_C,S,-1}, \phi_{T_{Ch},S,-1}$ | $\phi_{T_{Co},S,1}$ | — |
| $H_{0F}$ vs. $H_{AF}$ | $\phi_{T_C,S,1}$ | $\phi_{T_C,S,1}$ | — | — |

## 5 Discussion

This study had two proximate aims: to develop a minimal sufficient statistic for effects of interspecific interactions on community assembly, and to develop maximally sensitive (UMPU) hypothesis tests for detecting those effects:

The minimal sufficient statistic, **T**_*U*_, is the total number of species occurrences and the number of sites at which no species were observed. It is counterintuitive that this statistic summarizes all of the information in a presence absence matrix about interspe cific interactions, and that it does so with a maximal degree of data compression. To see this, consider the other statistics that have been developed on intuitive grounds: *T*_*C*_ and *T*_*Ch*_ summarize the extent to which species co-occur, which intuitively should decrease with effects of antagonistic interactions and increase with effects of facilitation (Stone and Roberts, 1990, Diamond, 1975). *T*_*Co*_ summarizes the diversity of communities,which should decrease with effects of ecological processes (communities should converge to a few stable states) (Pielou and Pielou, 1968). Last, *T*_*V*_ is an estimate of the covariance between the co-occurrence of different species, which again should decrease with antagonistic interactions (Schluter, 1984). However, Theorem 2 proves that while these statistics are reasonable, they discard key information about interspecific interactions under model *ℳ*. The counter intuitiveness of **T**_*U*_ illustrates the usefulness of the optimization approach for deriving predictions from hypothesized ecological phenomena.

The fact that the number of empty sites contains key information about effects of interspecific interactions potentially raises a need for new data. It is likely that biases exist against publishing data on sites at which no species were observed (Gotelli, 2000). Moreover, it is unclear whether sites where no species are observed are really devoid of species (MacKenzie et al, 2006). Additional studies are needed to address these matters.

Importantly, the sufficiency of **T**_*U*_ is contingent on the community assembling according to model *ℳ*. Hence, it is natural to ask whether *T*_*C*_, *T*_*Ch*_, *T*_*Co*_, and *T*_*V*_ are sufficient for other models. While this possibility requires further investigation, it appears not to be the case. One of the original aims of this study was to define these models, but despite extensive searching, none could be found.

Sufficient statistics are key not only for summarizing data, but also for deriving optimal hypothesis tests (Lehmann and Romano, 2005, Casella and Berger, 2002). The tests *ϕ*_*UC*_ and *ϕ*_*FC*_, which both use **T**_*U*_, are UMPU by Theorems 3 and 4 for testing for antagonistic interactions and facilitation under *ℳ*, respectively. The numerical analyses confirmed these results, and showed that under model *ℳ* these tests greatly outperform existing NMTPAs, correctly rejecting the null hypothesis at rates approaching twice that of existing tests. These tests also had the lowest Type I error rates. Given the importance of NMTPAs for addressing ecological questions and for formulating management policy, these improvements should be of practical importance.

In the foregoing analyses, randomized tests were used because otherwise comparable power functions could not have been constructed. However, for ecological applications, it is generally best to use nonrandomized analogues of *ϕ*_*UC*_ and *ϕ*_*FC*_ (i.e., versions in which *γ ≡* 0; see Definition 1) (Lehmann and Romano, 2005). The size of such tests should be confirmed numerically, as it may be much smaller than *α*. Interestingly, the results of *ϕ*_*UC*_,*ϕ*_*FC*_,andtheirnonrandomized analogues were identical for the two data sets analyzed here.

The UMP properties of *ϕ*_*UC*_ and *ϕ*_*FC*_ are proven under model *ℳ*, which assumes equivalence between species and a slow rate of immigration. These assumptions are particularly reasonable for communities that are spatially isolated; those composed of poorly dispersing species; and those located in homogeneous environments. However, for other data sets, *ℳ* may be inappropriate. The numerical results show that if deviations from *M* are small to moderate (e.g., model *ℳ*_1_, in which sites are assumed moderately heterogenous), then *ϕ*_*UC*_ and *ϕ*_*FC*_ can still perform well, and in some circumstances, optimally. This fact – that tests that are optimal under simplified models also tend to perform well under other models – provides a major justification for seeking optimal NMTPAs and other statistical methods under simplified models.

Despite the superior performance of *ϕ*_*UC*_ and *ϕ*_*FC*_, other NMTPAs have desirable properties which should not be overlooked.Although*ϕ*_*UC*_and*ϕ*_*FC*_ are robust to small to moderate deviations from *ℳ*, under different models, other NMTPAs may have greater power. Moreover, as noted above, the reliability of any hypothesis test depends on the inferences that are being drawn from it. It may be of interest to test other hypotheses than the ones examined here (*H*_0*C*_, *H*_*AC*_, *H*_0*F*_, and *H*_*AF*_), and in these instances other tests may perform better.

Another key consideration in implementing NMTPAs is robustness. A robust NMTPA has controlled Type I error rates under a broad range of models, or equivalently, under a generally realistic model (Lehmann and Romano, 2005). This consideration is important for NMTPAs because it is often difficult to verify assumptions, for instance, the unconditional probabilities of species occurring at each site (Ladau, 2008). Conditioning a test on both the row and column totals is one way to impart robustness: such tests are robust under models in which each species is equally likely to occur at all sites (site equivalency), or at each site, all species are equally likely to occur (species equivalency). These tests can also have substantial power, although their optimality is unclear (Ladau et al, in preparation). Additionally, (Ladau and Schwager, 2008) recently derived a NMTPA under a robustness constraint. This test requires information on the functional or taxonomic groups to which each species belongs, and can be proven UMP (Ladau and Schwager, unpublished results). In general, developing NMTPAs that are optimal within certain classes of tests is a promising area for further research.

In a general sense, these results demonstrate the feasibility of finding optimal null model tests in ecology. Optimality has heretofore not been considered explicitly in the development of null model tests. *A priori*, one might expect that ecological problems are too complicated to be amenable to the development of mathematically optimal methods. The results presented here show that the development of optimal null models tests is indeed possible, and they suggest that other complex inferential problems in ecology may yield to this approach.

This paper presents a new paradigm for developing optimal NMTPAs, other null model tests, and statistical methods in ecology. To develop these methods, a statistical model (set of candidate generating processes) should first be explicitly defined. Constraints should then be imposed on the method for instance, unbiasedness or robustness. With these constraints in place, attempts can then be made to solve the resulting optimization problem (Lehmann and Romano, 2005). This paradigm closely follows the approach to hypothesis testing introduced by Neyman and Pearson (Lehmann, 1993), an approach that has been used to great effect in the development of statistical methods in many fields.

## 6 Acknowledgements

This work was funded by the Santa Fe Institute and the Gordon and Betty Moore Foundation. Jennifer Dunne, Steven J. Schwager, Katherine S. Pollard, and Jon F. Wilkins provided helpful suggestions and insights.

## 7 Appendix

### Definitions

This section presents definitions that are necessary to prove the results in section 6.2. Some of the definitions are also presented heuristically in the text, but here they are made precise. Following (Ladau, 2008), the model used here is a set of probability measures on common sample space. Define *m* as the total number of species observed, *n* as the number of sites sampled, *M ≡ {*1, *…, m}*, and *N ≡ {*1, *…, n}*. The sample space is denoted by Ω: it consists of all possible presence absence matrices with *m* rows and *n* columns. Define 𝒫 (Ω) as the power set of Ω, and let *U* be the set of all probability measures on (Ω, *𝒫* (Ω)). For *i ∊ M* and *j ∊ N* define the random variables *X*_*ij*_ : Ω *→ {*0, 1*}* such that for any **b** ∊ Ω,

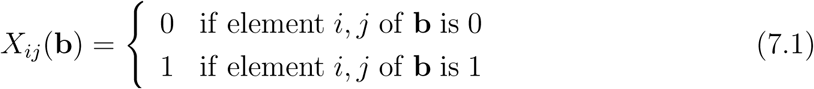

Hence, *X*_*ij*_ is an indicator variable for presence of species *i* at site *j*. For *i ∊ M* and *j ∊ N*, define *p*_*ij*_ : *𝒰 →* [0, 1] such that for *P ∊ 𝒰*, *p*_*ij*_(*P*) = *P* (*X*_*ij*_ = 1), and define *ξ* : *𝒰 →* [0, 1] such that *ξ*(*P*) = *P* (*X*_11_ = 1, *X*_21_ = 1)*/* [*P* (*X*_11_ = 1)*P* (*X*_21_ = 1)]. For convenience, let *p ≡ p*_11_. Let **C**_*j*_ : Ω *→* 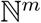 be a random vector such that for any **b** ∊ Ω, **C**_*j*_(**b**) gives the *j*th column of **b**. The model, *ℳ*, is the subset of *𝒰* such that for any *P ∊ ℳ* :

1. **C** _1_, *…,* **C**_*n*_ are mutually independent.
2. For any *j ∊ N* and **c** _1_, **c** _2_ ∊ N^*m*^: if *|***c**_1_*|*_1_=*|***c**_2_*|*_1_, then *P* (**C**_*j*_ = **c** _1_) = *P* (**C**_*j*_ = **c** _2_).
3. For all *i ∊ M* and *j ∊ N*, *p*_*ij*_ = *p*.
4. For any *j∊N* and nonempty *µ⊆M*, *P* 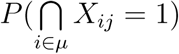 = *ξ*^*|µ|−*1^*p*^*|µ|*^.

Turning to statistics, for *i ∊ M*, define the random variables *U*_*i*_ : Ω *→ N* such that for **b** ∊ Ω, *U*_*i*_(**b**) gives the number of columns in **b** with sum *i*; i.e., the number of sites with exactly *i* species. Define the random variable *S* : Ω *→ {*0, 1, *…, nm}* so that for any presence absence matrix **b**, *S*(**b**) gives the total number of species occurrences in the matrix. Let **T**_*U*_ *≡* (*U*_0_, *S*) and *T*_*C*_, *T*_*Ch*_, *T*_*Co*_, *T*_*V*_ : Ω *→* ℝ denote the *C* score, number of checkerboards, number of species combinations, and *V* ratio statistics, respectively.

Some additional definitions are also necessary. For any set *E*, and nonnegative integer *k*, let *𝒫*_*k*_(*E*) denote the set of all *k* subsets of *E*. Given *j ∊ N*, define the function *A* : *𝒫*(*M*) *→ 𝒫*(Ω) such that for any *µ ⊆ M*,

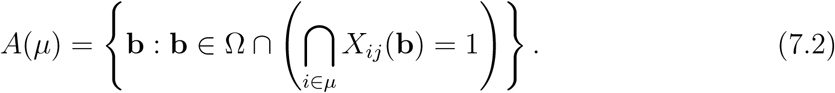

Likewise, define the function *Z* : *𝒫*(*M*) *→ 𝒫*(Ω) such that for any *µ ∊ 𝒫*(*M*),

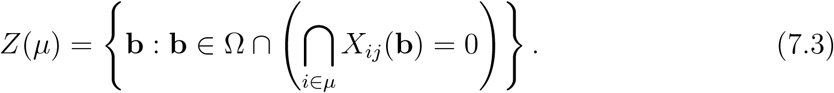

*A*(*µ*) and *Z*(*µ*) are the events that all and zero of the species in *µ* occur at site *j*, respectively. Define the random variables *S*_*j*_ : Ω *→ M* such that for any **b** ∊ Ω, *S*_*j*_(**b**) gives the sum of column *j* of **b**. For any *P∊ ℳ*, *s ∊ {*0, 1, *…, mn}*, and **b** ∊ Ω, let *P*^*s*^ denote the conditional probability of **b** given *S* = *s*; i.e., if *P*(*S* = *s*) *>* 0, then *P*^*s*^(**b**) = *P*(**b**, *S* = *s*)*/P*(*S* = *s*). Given *p*^*\**^ ∊ [0, 1], let 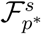 and let *f*_*S*_(*s*) = *P* (*S* = *s*); i.e., *f*_*S*_(*s*) is the mass function of *S*. Define the function space ϕ _*α*_ as the set of all size *α* tests on Ω having Neyman structure with respect to *S*.

### 7.1 Results

**Lemma 1.** *For any j ∊ N ; µ ⊆ M ; and distinct µ*_1_, *µ*_2_ *⊆ µ; A*(*µ*_1_) *∩ Z*(*µ − µ*_1_) *and A*(*µ*_2_) *∩ Z*(*µ − µ*_2_) *are disjoint.*

*Proof.* Given *j ∊ N*, *µ ⊆ M*, and distinct *µ*_1_,*µ*_2_ *⊆ µ*, the proof will follow by contradiction. Assume that there exists **b** ∊ Ω such that

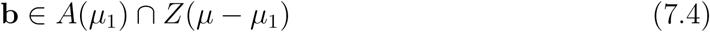

and

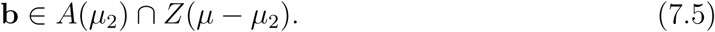

Because *µ*_1_ and *µ*_2_ are distinct, the symmetric difference of *µ*_1_ and *µ*_2_ is nonempty. Without loss of generality, fix *i ∊ µ*_1_ such that *i ∊ µ*_2_. The definition of *A* and (7.4) imply that *X*_*ij*_(**b**) = 1. However, because *µ*_1_ *⊆ µ*, *i ∊ µ*, so *i ∊ µ µ*_2_ and (7.5) implies that *X*_*ij*_(**b**) = 0. *X*_*ij*_(**b**) cannot simultaneously be 0 and 1, so the result follows.

**Lemma 2.** *For any j ∊ N,µ ⊆ M, and k ≤ |µ|*

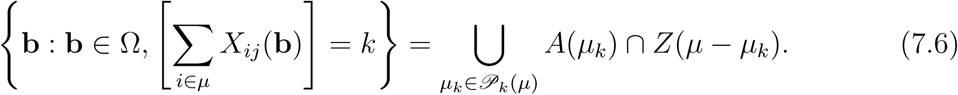

*Proof.* Given *j ∊ N*, *µ ⊆ M*, *k ≤ |µ|*, and **b**^*\**^ ∊ Ω, assume first that

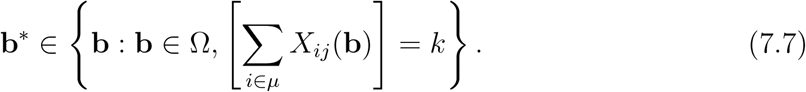

Equation (7.7) implies that there exists a subset *µ*_*k*_^*\**^ of *µ* with *k* elements such that *X*_*ij*_(**b**^*\**^) = 1 for all *i ∊ µ*_*k*_^*\**^, and *X*_*ij*_(**b**^*\**^) = 0 for all *i ∊ µ - µ*_*k*_^*\**^. Fix *µ*_*k*_^*\**^. Because *µ*_*k*_^*\**^ is a *k* subset of *µ*, it is an element of *𝒫*_*k*_(*µ*), implying that

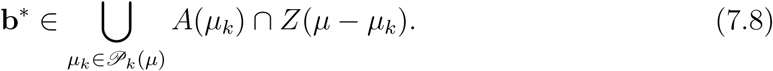

Begin now by assuming (7.8). By Lemma 1, the sets in the union in (7.8) are disjoint. It follows that there exists a *k* subset *µ*_*k*_^*\**^ ∊ *𝒫*_*k*_(*µ*) such that **b**^*\**^ ∊ *A*(*µ*_*k*_^*\**^) *2ȩ Z*(*µ - µ*_*k*_^*\**^). Fix *µ*_*k*_^*\**^. Because **b**^*\**^ ∊ *A*(*µ*_*k*_^*\**^) *∩ Z*(*µ - µ*_*k*_^*\**^), there are exactly *k* elements *i* in *µ*_*k*_^*\**^ such that *X*_*ij*_(**b**^*\**^) = 1. Equation (7.7) follows.

**Lemma 3.** *For any P∊ ℳ, j ∊ N, and* **b** ∊ Ω, *if S*_*j*_(**b**) *>* 0 *then*

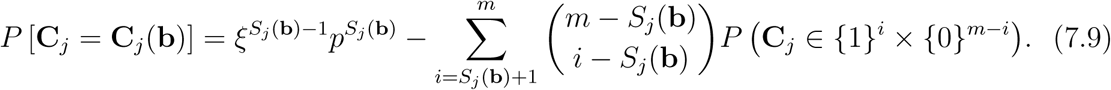

*Proof.* Given *P∊ ℳ*, *j ∊ N*, and **b** ∊ Ω, let *µ ≡* {*i* : *i ∊ M,* **C**_*j*_(**b**)(*i*) = 1 } and for convenience, let *s*_*j*_ *≡ S*_*j*_(**b**). Assume that *s*_*j*_ *>* 0. By basic probability,

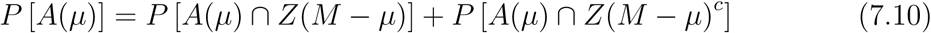

By definition, *A*(*µ*) *∩ Z*(*M - µ*) = { **C**_*j*_ = **C**_*j*_(**b**) }, and by Property 4 of *Mℳ*, *P*[*A*(*µ*)] = *ξ*^*s*^*j* 1*P*^*s*^*j*. Thus, it will suffice to show that

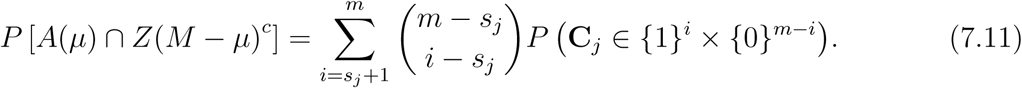

*Z*(*M - µ*) is the event that none of the species in *M - µ* occur at site *j*, so *Z*(*M - µ*)^*c*^ is the event that at least one of the species in *M -µ* occurs. Hence,

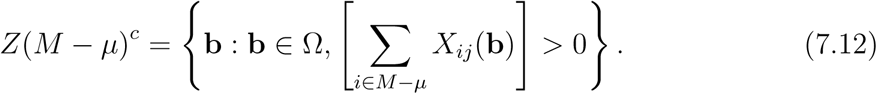

By definition, *|M - µ|* = *m - s*_*j*_, so this set is equivalent to

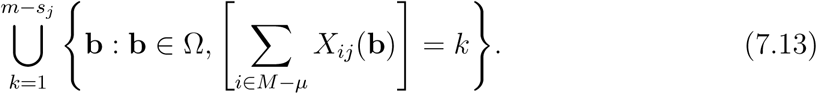

By Lemma 2,

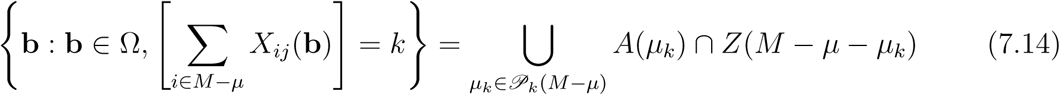

It follows from equations (7.12), (7.13), (7.14), and associativity that

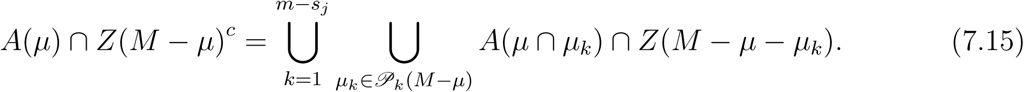

By Lemma 1, the sets *A*(*µ ∩ µ*_*k*_) *∩ Z*(*Mµ µ*_*k*_) in (7.15) are disjoint.Thus,

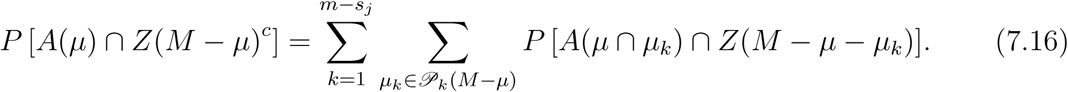

Property 2 of *ℳ* gives that for any *µ*_*k*_ *∊ 𝒫*_*k*_(*M - µ*),

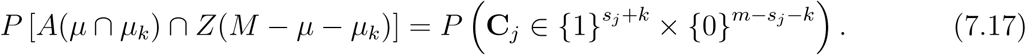

There are 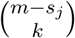 disjoint *k* subsets of *M - µ*, so

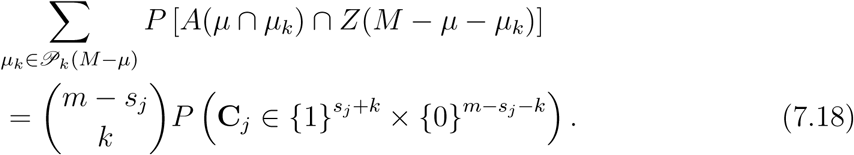

The proof is completed by substituting (7.18) into (7.16) and defining *i ≡ k* + *s*_*j*_.

**Lemma 4.** *For any P∊ ℳ, j ∊ N, and* **b** ∊ Ω, *if S*_*j*_(**b**) *>* 0:

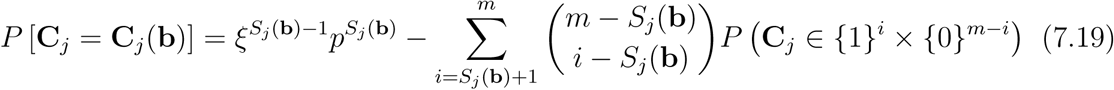

*if and only if*

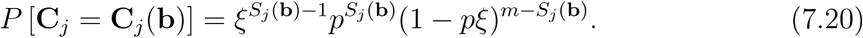

*Proof.* Given *P∊ ℳ*, *j ∊ N*, and **b** ∊ Ω, assume *S*_*j*_(**b**) *>* 0. The proof of the implication from (7.19) to (7.20) will follow by reverse induction on *S*_*j*_(**b**). If *S*_*j*_(**b**) = *m*, then the result follows trivially. Assume now that for some integer 2 *≤ k ≤ m*, the result holds if *S*_*j*_(**b**) *≥ k*, and that *S*_*j*_(**b**) = *k −* 1. By (7.19) and the induction hypothesis,

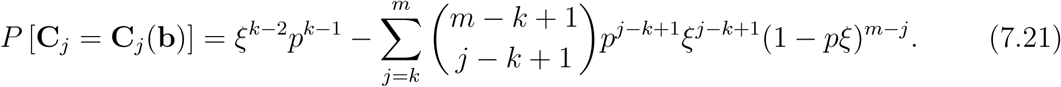

Defining *i ≡ j - k* + 1, the right hand side of (7.21) is algebraically equivalent to

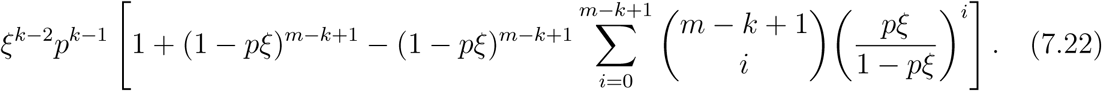

By the binomial theorem,

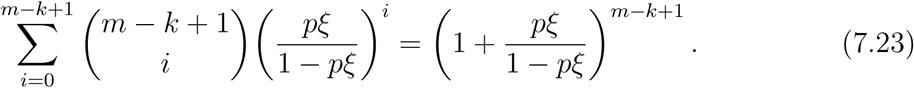

Substituting (7.23) into (7.22) and simplifying yields, by (7.21),

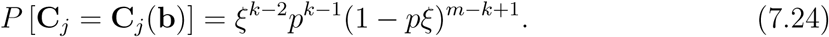

Because *S*_*j*_(**b**) = *k −* 1, the implication from (7.19) to (7.20) is proven. The arguments can be reversed to show the converse.

**Lemma 5.** *For any P∊ ℳ and* **b** ∊ Ω,

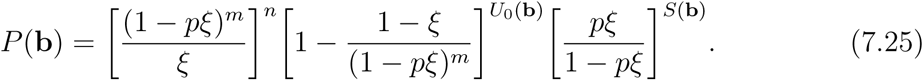

*Proof.* Given *P∊ ℳ* and **b** ∊ Ω, by property 1 of *ℳ*,

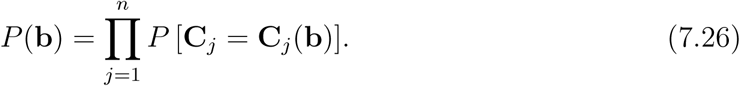

For convenience, define *v ≡* {*j* : *j ∊ N, S*_*j*_(**b**) *>* 0 }. Applying Lemmas 3 and 4,

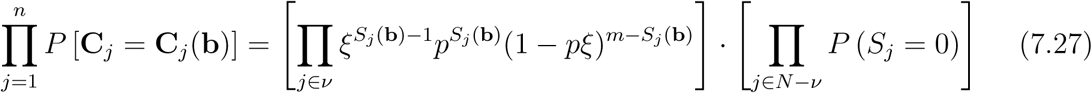

Examining the first term in (7.27), *|v|* = *n - U*_0_(**b**) and 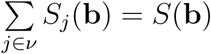 so

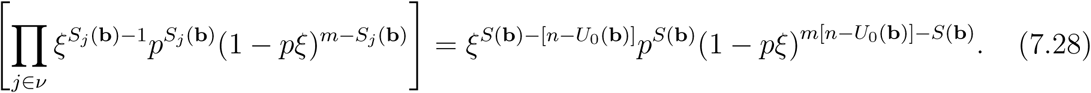

Turning to the second term in (7.27), for any *j ∊ N*,

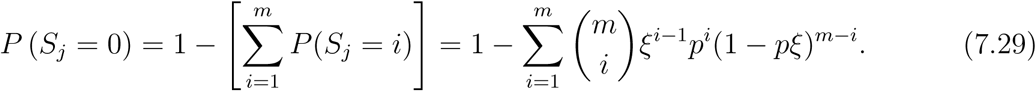

By elementary algebra,

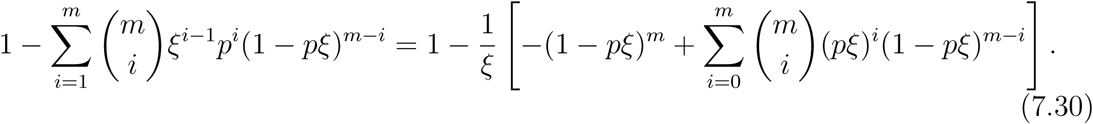

Applying the binomial theorem and simplifying yields

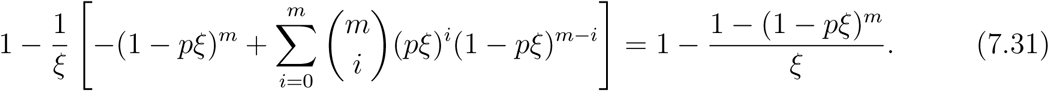

Because *|N - v|* = *U*_0_(**b**),

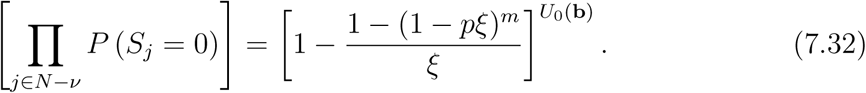

Substituting (7.28) and (7.32) into (7.27) and simplifying yields the result.

**Lemma 6.** *For any* **b** _1_, **b** _2_ ∊ Ω, *P*(**b** _1_)*/P*(**b** _2_) *is constant as a function of ξ and Pif and only if* **T**_*U*_ (**b** _1_) = **T**_*U*_ (**b** _2_).

*Proof.* Given **b** _1_, **b** _2_ ∊ Ω, for convenience, let *d*_*U*_ *≡ U*_0_(**b** _1_) *- U*_0_(**b** _2_) and *d*_*S*_ ≡ *S*(**b** _1_) *-S*(**b** _2_). It follows that from Lemma 5 and (7.25) that

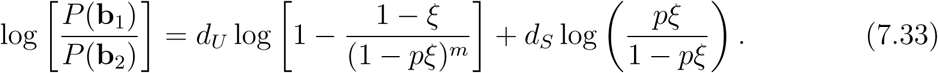

Clearly, the right hand side of (7.33) is not a function of *ξ* and *P* if and only if *d*_*U*_ = 0 and *d*_*S*_ = 0; i.e., **T**_*U*_ (**b** _1_) = **T**_*U*_ (**b** _2_).

**Theorem 1.** *The statistic* **T**_*U*_ *is minimal sufficient for ℳ*

*Proof.* The sufficiency of **T**_*U*_ follows immediately from applying the Factorization Criterion (Casella and Berger, 2002) to (7.25). Minimal sufficiency follows from Lemma 6 and applying Theorem 6.2.13 of (Casella and Berger, 2002).

**Theorem 2.** *The statistics T*_*C*_, *T*_*Ch*_, *T*_*Co*_, *and T*_*V*_ *are not sufficient for ℳ.*

*Proof.* If the four statistics are sufficient, then by Theorem 1 and the definition of minimal sufficiency, for all **b** ∊ Ω, **T**_*U*_ (**b**) is a function of these statistics. Hence, to prove the result, for each statistic it will suffice to find **b**^*\**^, **b**^*\*\**^ ∊ Ω such that *T*_*i*_(**b**^*\**^) = *T*_*i*_(**b**^*\*\**^) for *T*_*i*_ ∊ {*T*_*C*_, *T*_*Ch*_, *T*_*Co*_, *T*_*V*_} but **T**_*U*_ (**b**^*\**^) *≠* **T**_*U*_ (**b**^*\*\**^). Define

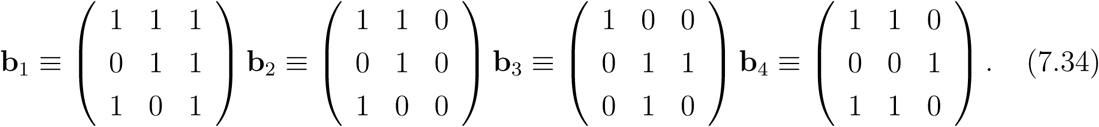

The values of each statistic for **b** _1_ to **b** _4_ are listed in Table 2. The result follows by noting that **T**_*U*_ differs for each value of **b**, but *T*_*C*_(**b** _1_) = *T*_*C*_(**b** _2_), *T*_*Ch*_(**b** _3_) = *T*_*Ch*_(**b** _4_), *T*_*Co*_(**b** _2_) = *T*_*Co*_(**b** _3_), and *T*_*V*_ (**b** _3_) = *T*_*V*_ (**b** _4_). Other counterexamples can be generated easily.

**Lemma 7.** *For any P*∊ [0, 1] *and ξ >* 0, *if* (1 *- 𝒫ξ*)^*m*^ - (1 *- ξ*) *>* 0, *then* (1 *- pξ*) *-mp*(1 *- ξ*) *≥* 0.

**Table 2:** Values of the statistics for **b**_1_ to **b**_4_.

| Matrix | Statistic |  |  |  |  |
| --- | --- | --- | --- | --- | --- |
| | $\mathbf{T}_U$ | $T_C$ | $T_{Ch}$ | $T_{Co}$ | $T_V$ |
| $\mathbf{b}_1$ | (0, 7) | 1/3 | 0 | 3 | 1/2 |
| $\mathbf{b}_2$ | (1, 4) | 1/3 | 1 | 3 | 4/3 |
| $\mathbf{b}_3$ | (0, 4) | 1 | 2 | 3 | 1/3 |
| $\mathbf{b}_4$ | (0, 5) | 4/3 | 2 | 2 | 1/3 |

*Proof.* Given *p* ∊ [0, 1] and *ξ >* 0, assume that (1 *- pξ*)^*m*^ - (1 *- ξ*) *>* 0. The proof will follow by a case analysis on *ξ*.

Case 1: If *ξ* = 1, then (1 *- pξ*) *- mp*(1 *- ξ*) = 1 *- p ≥* 0.

Case 2: *ξ >* 1. Because (1 *- pξ*) *- mp*(1 *- ξ*) is nonnegative when *ξ* = 1 and

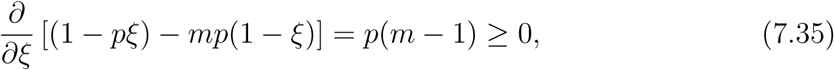

the result follows.

Case 3: *ξ <* 1. For convenience, define *r ≡* (*m −* 1)*/m*. By the Bernoulli inequality (Mitrinovic, 1970), 1 *- ξr >* (1 *- ξ*)^*r*^. It follows from *ξ <* 1 that

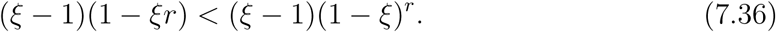

Algebraic manipulation of (7.36) yields

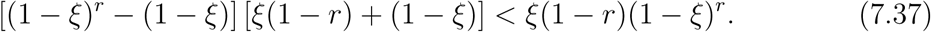

As *ξ <* 1 and 1 *- r* = 1 *−* (*m −* 1)*/m >* 0, (7.37) implies that

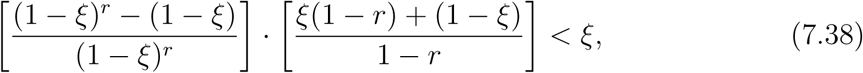

so

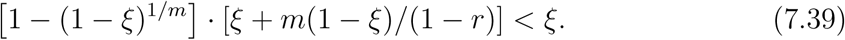

Because *ξ <* 1, (7.39) can be written as:

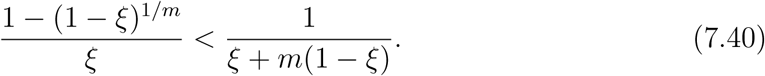

Last, by assumption (1 *- pξ*)^*m*^ - (1 *- ξ*) *>* 0, so *p <* [1 *−* (1 *- ξ*)^1/*m*^]*/ξ*, implying by (7.39) that

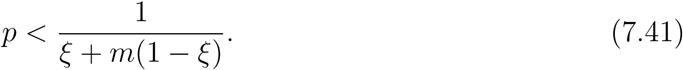

Algebraic manipulation of (7.41) yields (1 *- pξ*) *- mp*(1 *- ξ*) *>* 0.

**Lemma 8.** *For any s*∊{0, 1, *…, mn*} and *p*^*\**^ and 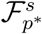 *has a monotone likelihood ratio.*

*Proof.* Given *s ∊* {0, 1, *…, mn*} and *p*^*\**^ ∊ [0, 1], by Lemma 5, 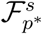 solely by *ξ*. Hence, it will be sufficient to show that for any 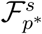 is indexed by the and **b** _1_, **b** _2_ ∊ Ω, if *S*(**b** _1_) = *S*(**b** _2_) = *s* and *U*_0_(**b** _1_) *> U*_0_(**b** _2_), then

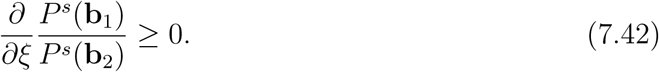

Let *d*_*U*_ *≡ U*_0_(**b** _1_) *- U*_0_(**b** _2_). By Lemma 5,

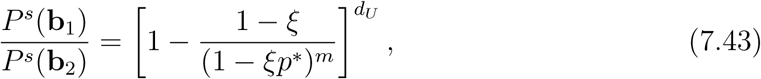

so

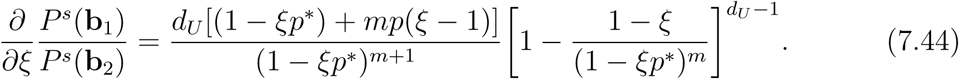

If *ξ* = 0, then (7.44) simplifies to *d*_*U*_ (1 *- mp*)(0)*/*(1), so the result follows. If *ξ >* 0, then Lemma 4 implies that

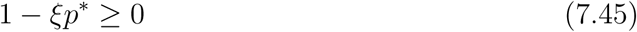

because otherwise *∃***c** *∊* {0, 1 }^*m*^ such that *P* [**C**_*j*_ = **c**] *<* 0. Moreover, equations (7.29) to (7.31) imply by similar reasoning that

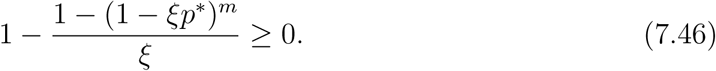

If (7.46) is identically 0, then 1 *−* (1 *- ξ*)*/*(1 *- ξp*^*\**^)^*m*^ = 0, and the result follows. Assuming that strict inequality holds for (7.46), by Lemma 7, (1 *- ξp*^*\**^) + *mp*(*ξ −* 1) *>* 0. Moreover, 1 *−* (1 *- ξ*)*/*(1 *- ξp*^*\**^)^*m*^ *>* 0, so by (7.45), equation (7.44) is nonnegative.

**Lemma 9.** *For any s ∊* {0, 1, *…, mn} and p*^*\**^ ∊ [0, 1], *U*_0_ *is sufficient for 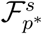. Proof.* Given *s ∊* {0, 1, *…, mn*}, *p*^*\**^ ∊ [0, 1], and 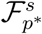, by Lemma 5, for any **b***\** ∊ Ω,

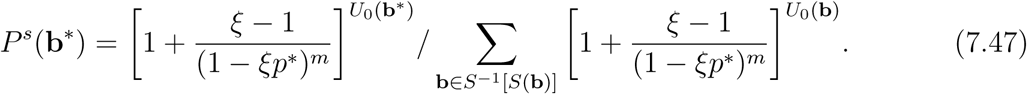

Equation (7.47) is a function only of **b**^*\**^ through *U*_0_, so the result follows from the factorization criterion (Casella and Berger, 2002).

**Lemma 10.** *For any s ∊* {0, 1, *…, mn*} *and p^*^* ∊ [0, 1], *if 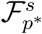, then ϕ*_*UC*_ *is a UMP size α test for testing H*_0*C*_ *versus H*_*AC*_ *conditional on S* = *s.*

*Proof.* For any *s ∊* {0, 1, *…, mn*} and *p*^*\**^ ∊ [0, 1], assume that *P^s^* ∊ 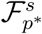. Ladau et al (in preparation) establishes that *ϕ*_*UC*_ is size *α*. Moreover, by Lemma 8, 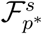 has an MLR, and by Lemma 9, *U*_0_ is sufficient for *Fs*. The Karlin Rubin Theorem (Lehmann and Romano, 2005, Casella and Berger, 2002) applies to yield the result.

**Lemma 11.** *ϕ*_*UC*_ *is a UMP size α test among ϕ* ∊ ϕ _*α*_.

*Proof.* Ladau et al (in preparation) establishes that *ϕ*_*UC*_ has Neyman structure with respect to *S* and has size *α*, making it an element of ϕ _*α*_. To establish that it has maximal power, for any *P ∊ ℳ* with *ξ ∊ H*_*AC*_, it is necessary to show that over *ϕ* ∊ ϕ _*α*_, *ϕ*_*UC*_ maximizes *E*_*P*_ *ϕ*(**b**). By definition,

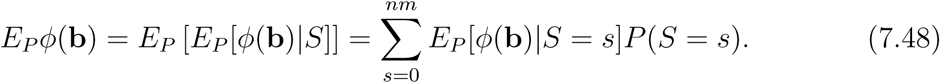

Lemma 10 implies that for each *s ∊* {0, 1, *…, mn*}, *ϕ*_*UC*_ maximizes *E*_*P*_ [*ϕ*(**b**)*|S* = *s*]. Thus, it maximizes the right hand side of (7.48) and the lemma follows.

**Theorem 3.** *ϕ*_*UC*_ *is a UMPU size α test of H*_0*C*_ *versus H*_*AC*_.

*Proof.* By Lemma 11, and Lemma 4.1.1 and Theorems 4.3.2 and 4.4.1 of (Lehmann and Romano, 2005), it will be enough to show that if *ξ* = 1, then *S* is sufficient for *𝒢 ≡* {*P : P ∊ ℳ, ξ* = 1} and *𝒢s;S ≡* {*f _S_* :*#x03BE;*(*P*) = 1 } is complete. To show sufficiency, by Lemma 5, for any *P ∊ 𝒢* and **b** ∊ Ω,

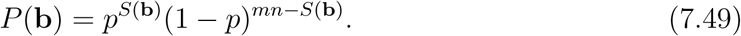

The factorization criterion (Casella and Berger, 200) applies to yield the result. To show completeness, for *s ∊* {0, 1, *…, mn*}, there are 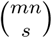 matrices **b** ∊ Ω such that *S*(**b**) = *s*, implying by (7.49) that for *f*_*S*_ *∊ 𝒢*^*S*^

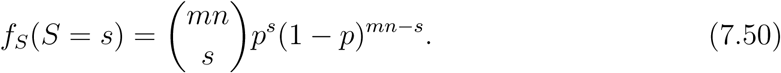

Thus, *S* is distributed binomial(*nm, p*). The binomial family is complete, implying the theorem.

**Theorem 4.** *ϕ*_*UF*_ *is a UMPU size α test of H*_0*F*_ *versus H*_*AF*_.

*Proof.* This result follows by arguments analogous to those of the proof of Theorem 3.

## References

Alatalo R (1982) Bird species distributions in the galapagos and other archipelagos: competition or chance? Ecology 63:881–887

Atmar W, Patterson B (1993) The measure of order and disorder in the distribution of species in fragmented habitat. Oecologia 96:373–382

Brown J, Kelt D, Fox BJ (2002) Assembly rules and competition in desert rodents. The American Naturalist 160:815–818

Casella G, Berger R (2002) Statistical Inference: Second Edition. Duxbury, Pacific Grove

Connor E, Simberloff D (1978) Species number and compositional similarity of the galapagos flora and avifauna. Ecological 48:219–248

Connor E, Simberloff D (1979) The assembly of species communities: Chance or competition? Ecology 60:1132–1140

Connor E, Simberloff D (1986) Competition, scientific method, and null models in ecology. American Scientist 74:155–162

Dayton G, Fitzgerald L (2001) Competition, predation, and the distributions of four desert anurans. Oecologia 129:430–435

Diamond J (1975) Assembly of species communities. In: Ecology and Evolution of Communities, Harvard University Press, pp 342–344

Dufrene M, Legendre P (1997) Species assemblages and indicator species: The need for a flexible asymmetrical approach. Ecological Monographs 67:345–366

Feeley K (2003) Analysis of avian communities in lake guri, venezuela, using multiple assembly rule models. Oecologia 137:104–113

Fox B, Brown J (1993) Assembly rules for functional groups in north american desert rodent communities. Oikos 67:358–370

Fox B, Brown J (1995) Reaffirming the validity of the assembly rule for functional groups or guilds: a reply to wilson. Oikos 73:125–132

Gilpin M, Diamond J (1982) Factors contributing to non-randomness in species co-occurrences on islands. Oecologia 52:75–84

Gotelli N (2000) Null model analysis of species co occurrence patterns. Ecology 81:2606–2621

Gotelli N (2001) Research frontiers in null model analysis. Global Ecology and Biogeography 10:337–343

Gotelli N, Abele L (1982) Statistical distributions of west indian land bird families. Journal of Biogeography 9:421–435

Gotelli N, Ellison A (2002) Assembly rules for new england ant assemblages. Oikos 99:591–599

Gotelli N, Graves G (1996) Null Models in Ecology. Smithsonian Institution, Washington

Gotelli N, McCabe D (2002) Species co-occurrence: a meta analysis of j. m. diamond's assembly rules model. Ecology 83:2091–2096

Gotelli N, Rohde K (2002) Co-occurrence of ectoparasites of marine fishes: a null model analysis. Ecology Letters 5:86–94

Gotelli N, Buckley N, Wiens J (1997) Co-occurrence of australian land birds: Diamond's assembly rules revisited. Oikos 80:311–324

Graves G, Gotelli N (1993) Assembly of avian mixed species flocks in amazonia. Proceedings of the National Academy of Science USA 90:1388–1391

Heino J, Soininen J (2005) Assembly rules and community models for unicellular organisms: patterns in diatoms of boreal streams. Freshwater Biology 50:567–577

Hubbell S (2001) The Unified Neutral Theory of Biodiversity and Biogeography. Princeton University Press, New Jersey

Ladau J (2008) Validation of null model tests using neyman pearson hypothesis testing theory. Theoretical Ecology 1:241–248

Ladau J, Schwager S (2008) Robust hypothesis tests for independence in community assembly. Journal of Mathematical Biology 57:537–555

Lehmann E (1993) The fisher, neyman-pearson theories of testing hypotheses: One theory or two? Journal of the American Statistical Association 88:1242–1249

Lehmann E, Casella G (1998) Theory of Point Estimation. Springer-Verlag, New York

Lehmann E, Romano J (2005) Testing Statistical Hypotheses: Third Edition. Springer, New York

MacKenzie D, Nichols J, Pollock K (2006) Occupancy Estimation and Modeling: Inferring Patterns and Dynamics of Species Occurrence. Elsevier, Amsterdam

McNab B (1971) The structure of tropical bat faunas. Ecology 52:352–358

Miklos I, Podani J (2004) Randomization of presence-absence mtarices: comments and new algorithms. Ecology 85:86–92

Mitchell-Olds T (2001) Arabidopsis thaliana and its wild relatives: a model system for ecology and evolution. Trends in Ecology and Evolution 16:693–700

Mitrinovic D (1970) Analytic Inqualities. Springer-Verlag, New York

Mouillot D, George Nascimento M, Poulin R (2005) Richness, structure and functioning in metazoan parasite communities. Oikos 109:447–460

Pielou D, Pielou E (1968) Association among species of infrequent occurrence: the insect and spider fauna of polyporus betulinus (bulliard) fries. Journal of Theoretical Biology 21:202–216

Reed T (1980) Turnover frequency in island birds. Journal of Biogeography 7:329–335

Roberts A, Stone L (1990) Island-sharing by archipelago species. Oecologia 83:560–567

Ross S (2006) Simulation: Fourth Edition. Academic Press, San Diego

Sanders N, Gotelli N, Heller N, Gordon D (2003) Community disassembly by an invasive species. Proceedings of the National Academy of Science USA 100:2474–2477

Schluter D (1984) A variance test for detecting species associations, with some example applications. Ecology 65:998–1005

Schluter D (1990) Species-for-species matching. American Naturalist 136:560–568

Simberloff D, Connor E (1981) Missing species combinations. American Naturalist 118:215–239

Sokal R, Rohlf F (1995) Biometry: the principles and practice of statistics in biological research. W.H. Freeman, New York

Stephens C, Heau J, Gonzalez-Rosas C, Ibarra-Cerdena C, Sanchez-Cordero V (2008) Using biotic interaction networks for prediction in biodiversity and emerging diseases. Nature Proceedings NA:preprint

Stone L, Roberts A (1990) The checkerboard score and species distributions. Oecologia 85:74–79

Stone L, Dayan T, Simberloff D (1996) Community-wide assembly patterns unmasked: The importance of species' differing geographical ranges. The American Naturalist 148:997–1015

Stone L, Dayan T, Simberloff D (2000) On desert rodents, favored states, and unresolved issues: Scaling up and down regional assemblages and local communities. The American Naturalist 156:322–328

Wilson J (1989) A null model of guild proportionality, applied to stratification of a new zealand temperate forest. Oecologia 80:263–267

